# Investigating dual Ca^2+^ modulation of the ryanodine receptor 1 by molecular dynamics simulation

**DOI:** 10.1101/736934

**Authors:** Wenjun Zheng, Han Wen

## Abstract

The ryanodine receptors (RyR) are essential to calcium signaling in striated muscles. A deep understanding of the complex Ca^2+^-activation/inhibition mechanism of RyRs requires detailed structural and dynamic information for RyRs in different functional states (e.g., with Ca^2+^ bound to activating or inhibitory sites). Recently, high-resolution structures of the RyR isoform 1 (RyR1) were solved by cryo-electron microscopy, revealing the location of a Ca^2+^ binding site for activation. Toward elucidating the Ca^2+^-modulation mechanism of RyR1, we performed extensive molecular dynamics simulation of the core RyR1 structure in the presence and absence of bound and solvent Ca^2+^ (total simulation time is > 5 microseconds). In the presence of solvent Ca^2+^, Ca^2+^ binding to the activating site enhanced dynamics of RyR1 with higher inter-subunit flexibility, asymmetric inter-subunit motions, outward domain motions and partial pore dilation, which may prime RyR1 for subsequent channel opening. In contrast, the solvent Ca^2+^ alone reduced dynamics of RyR1 and led to inward domain motions and pore contraction, which may cause inhibition. Combining our simulation with the map of disease mutation sites in RyR1, we constructed a wiring diagram of key domains coupled via specific hydrogen bonds involving the mutation sites, some of which were modulated by Ca^2+^ binding. The rich structural and dynamic information gained from this study will guide future mutational and functional studies of RyR1 activation and inhibition by Ca^2+^.

**Statement of Significance:** The ryanodine receptors (RyR) are key players in calcium signaling, and make prominent targets for drug design owning to their association with many diseases of cardiac and skeletal muscles. However, the molecular mechanism of their activation and inhibition by Ca^2+^ remains elusive for the lack of high-resolution structural and dynamic information. Recent solutions of RyR1 structures by cryo-EM have paved the way for structure-based investigation of this important receptor by atomistic molecular simulation. This study presented, to our knowledge, the most extensive MD simulation of RyR1 core structure. Our simulation has offered new insights to the dual modulation mechanism of Ca^2+^, in which Ca^2+^ binding to the activating site primes RyR1 activation by elevating its dynamics while solvent Ca^2+^ inhibits RyR1 by reducing its dynamics. Additionally, our simulation has yielded a new wiring diagram of the allosterically coupled RyR1 domains informed by disease mutations.

## Introduction

Calcium (Ca^2+^) signaling is critically involved in many physiological processes including the excitation-contraction coupling in skeletal and cardiac muscles. As a key Ca^2+^ channel that regulates Ca^2+^ concentration, ryanodine receptors (RyRs) (1, 2) undergo a closed-to-open activation transition to release Ca^2+^ from the sarcoplasmic reticulum in response to an action potential that activates the voltage-gated calcium channels which subsequently activate RyRs (3). RyRs are activated by micromolar Ca^2+^ and inhibited by millimolar Ca^2+^ (4–8) via different Ca^2+^ binding sites (i.e. activating and inhibitory sites) accessible to both cytosolic and luminal Ca^2+^ (9) (10, 11). RyRs are also subject to regulations by various agents including ATP, Mg^2+^, phosphorylation, and via interactions with other proteins such as calmodulin (12), which involve various functional domains of RyRs (13). Remarkably, the binding sites for various agents are separated from the channel pore by as much as > 200 Å in RyR1 (2), highlighting the importance of allosteric couplings to RyR1 activation.

In three isoforms of RyRs (RyR1, RyR2, and RyR3), more than 500 mutations have been linked to human diseases including malignant hyperthermia, central core disease, and various heart disorders (14–16), making RyRs a prominent target for drug design (17). Many of these mutations cause excess RyRs activity, possibly by perturbing inter-domain/subunit interactions that favor the closed state of the channel (18). For the lack of high-resolution structures, the detailed mechanisms for the activation/inhibition of RyRs remain unknown. Indeed, it was highly challenging to solve the full-length structure of RyRs, owning to their enormous size (~2.2 MDa and >20000 residues) and high flexibility. Thanks to the recent revolution in cryo-electron microscopy (EM), high-resolution (≥3.6Å) structures of RyR1 were finally solved in different forms (e.g. the apo closed form, the Ca^2+^-bound primed form, and the Ca^2+^/ATP/Caffeine-bound open form) (19–24). Together, these new structures offered unprecedented insights to the global architecture and conformational changes in RyR1, and enabled our previous atomistic simulation of the N-terminal domains of RyR1 (25) and coarse-grained modeling of the closed-to-open transition of RyR1 with amino-acid level of details (26).

The structural architecture of RyR1 features a homo-tetramer of four subunits (see Fig 1), each containing a large cytoplasmic moiety (~80% of the total mass) and a trans-membrane domain (TMD). The cytoplasmic moiety is comprised of an N-terminal domain (NTD) and a large α-solenoid domain, surrounded by a number of peripheral domains including three SPRY domains and two tandems of repeat domains, and a core α-solenoid domain (CSOL) including a pair of EF-hand motifs (EFH12). A recent study showed that this EF-hand Ca^2+^ binding domain is important for luminal Ca^2+^ activation of RyR2 but not essential for cytosolic Ca^2+^ activation of RyR2 (27). The NTD and CSOL are rich in disease mutations (14). At the CSOL-TMD interface is a thumb and forefingers domain (TAF). The TMD includes six transmembrane helices (S1-S6) reminiscent of the voltage-gated ion channel superfamily. The S1-S4 helices form the pseudo voltage sensor domain (PVSD). The S5 and S6 helices form a pore domain that encloses a central pore for Ca^2+^ flow. The PVSD and pore domain are coupled via a S4-S5 linker (S4S5L), which provides a functionally critical interface for RyR channel gating (28). The TMD has two unique features at its cytoplasmic interface, including a cytoplasmic extension of S6 helix (S6C) capped by a C-terminal domain (CTD), and a helical bundle subdomain between S2 and S3 helices (S2S3). The CTD harbors a Ca^2+^-binding site (22, 29) critical for activation, including residues E3893, H3895, E3967, Q3970, and T5001 at the CSOL/CTD interface. The S2S3 forms inter-subunit contacts with the Ca^2+^-binding EFH12. The strategic locations of these domains make them promising candidates for allosterically coupling Ca^2+^ binding to the channel pore (19–22). While the static cryo-EM structures (19–22) hint for possible allosteric coupling pathways for RyR1 activation, the functional significance of such hypothetical pathways must be tested by directly probing the dynamics of RyR1 activation under physiological conditions.

**Figure 1.**
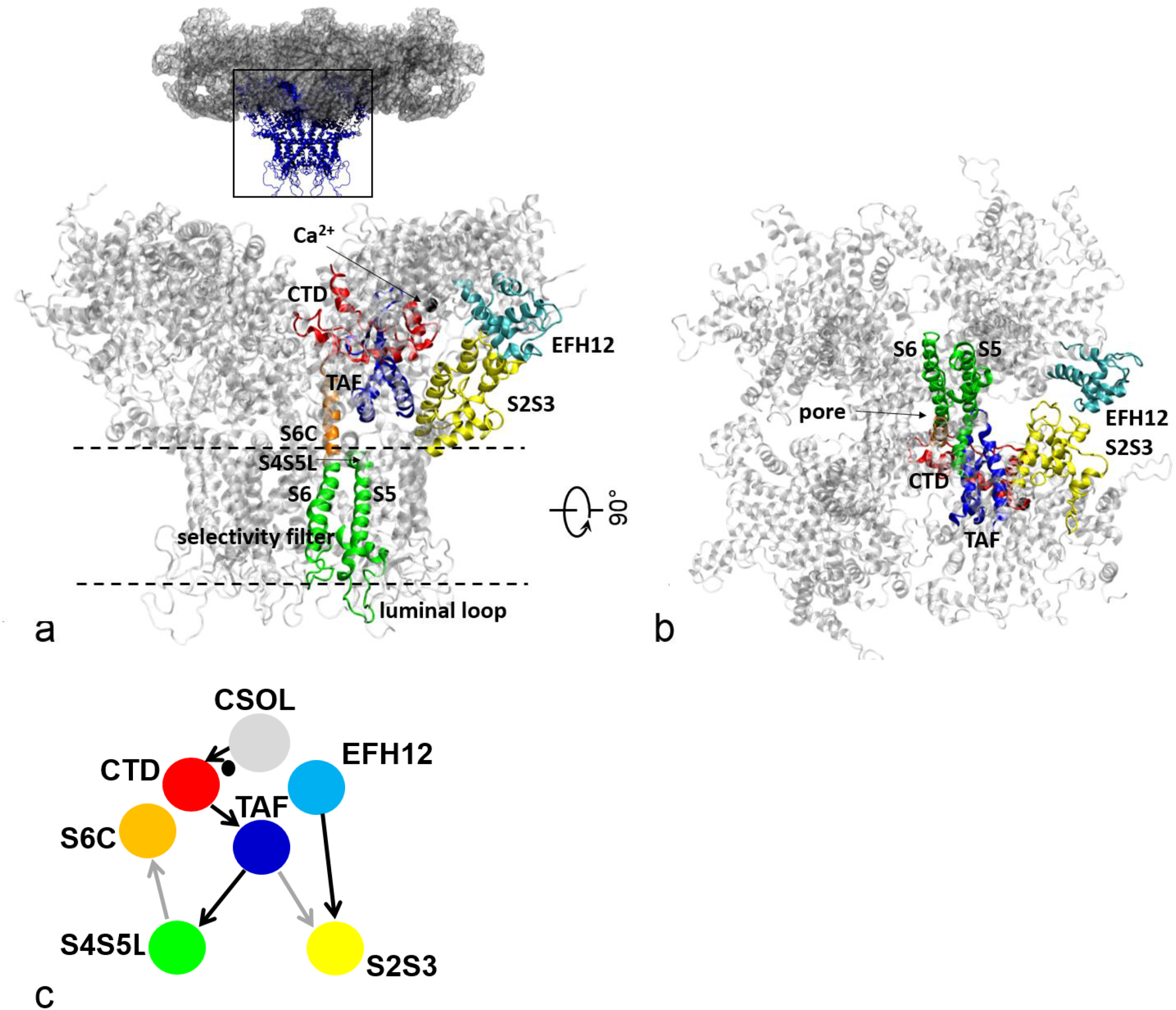
The architecture of the RyR1 core structure: (a) in side view; (b) in bottom view. The key domains of one representative subunit are colored as follows: CTD (red), TAF (blue), S6C (orange), pore domain (green), S2S3 (yellow), and EFH12 from an adjacent subunit (cyan). At the top of panel a is a full-length RyR1 with the core structure boxed and colored in blue. In panel a, the two parallel dash lines mark the transmembrane domains of the RyR1 core. Panel c shows the wiring diagram of key domains deduced from our mutation sites based hydrogen bond analysis, where Ca^2+^-dependent couplings are indicated by black arrows, and Ca^2+^-independent couplings are indicated by gray arrows.

Molecular Dynamics (MD) simulation is the method of choice for exploring protein dynamics and energetics under physiological conditions at atomic resolution (30), including the pore domain of RyR1 (31, 32). In the past, for a large biomolecular system with explicit solvent, MD simulation was highly expensive. Thanks to recent developments in computing hardware and software, particularly the use of graphics processing units to accelerate MD simulation (33), one can now routinely run MD simulation of a solvated protein (with several hundred thousand atoms) at a speed of several nanoseconds per day on a single computer node. This enables effective sampling of the conformational space by simultaneously running multiple relatively short MD trajectories (~ hundreds of nanoseconds) (34). Such simulation strategy is suitable for probing energetics and fast dynamics of a protein complex in a stable state, although much longer MD simulations (microseconds to milliseconds) are needed to explore slow dynamics such as the activation transition in RyRs.

Toward elucidating the Ca^2+^-modulation mechanism of RyR1, we performed extensive MD simulation of the core RyR1 structure in the presence and absence of bound and solvent Ca^2+^. Our simulation focused on the C-terminal core structure of RyR1 (see Fig 1), which was best resolved by cryo-EM (22). Our findings supported dual effects of Ca^2+^ modulation of RyR1: 1. Ca^2+^ binding (at the activating site) elevated dynamics of RyR1 with higher inter-subunit flexibility, asymmetric inter-subunit motions, outward domain motions, and partial pore dilation, which may prime RyR1 for subsequent channel opening; 2. Solvent Ca^2+^ alone reduced RyR1 dynamics and led to channel contraction, which may drive RyR1 inhibition. Combining our simulation with the map of disease mutation sites in RyR1, we have constructed a wiring diagram of key domains linked by specific hydrogen bonds involving the mutation sites, hinting for possible allosteric coupling pathways (through the CTD and TAF domains) for Ca^2+^ activation of RyR1. The structural and dynamic insights gained from this study will guide future mutational and functional studies of RyR1 activation and inhibition by Ca^2+^.

## Materials and Methods

### MD simulation setup

We performed MD simulation for the core structure (residues 3640 – 5037) of an RyR1 tetramer truncated from a 3.6-Å cryo-EM structure of Ca^2+^-bound rabbit RyR1 (PDB id: 5T15), which is the best-resolved RyR1 structure published to date. This structure is thought to capture a key intermediate state between the closed state and the open state in which the receptor is “primed” by Ca^2+^ binding for subsequent activation (22). The pre-oriented Ca^2+^-bound RyR1 structure was retrieved from the OMP database (https://opm.phar.umich.edu/). We used the MODLOOP server (35) to add two short missing loops (residues 3742 - 3746 and 4588 - 4625), while another long missing segment (residues 4254 - 4539) were not modeled. We then used the Membrane Builder module (36–38) of the CHARMM-GUI webserver (39, 40) to embed the RyR1 core structure (with or without four bound Ca^2+^) into a bilayer of 1-palmitoyl-2-oleoyl phosphatidylcholine (POPC) lipids surrounded by a box of water molecules, Ca^2+^ or K^+^, and Cl^−^ at a concentration of 0.15 M, with a 15-Å buffer of water/lipids extending from the protein in each direction. Each system has a total of ~638500 atoms. After energy minimization, six steps of equilibration were performed (with gradually reduced harmonic restraints applied to protein, lipids, water, and ions). Finally, we conducted production MD runs in the NPT ensemble. The Nosé-Hoover method (41, 42) was used with a temperature of 303 K. The Parrinello–Rahman method (43) was used for pressure coupling. A 10-Å switching distance and a 12-Å cutoff distance were used for nonbonded interactions. The particle mesh Ewald method (44) was used for electrostatics calculations. The LINCS algorithm (45) was used to constrain the hydrogen-containing bond lengths, which allowed a 2-fs time step for MD simulation. The energy minimization and MD simulation were performed using the GROMACS program (46) version 5.0.3-gpu, the CHARMM36 force field (47, 48), and TIP3P water model (49). For each system, nine 200-ns MD trajectories were generated and combined for further analysis (after discarding the initial 50 ns for system equilibration).

### RMSF analysis

To quantify the per-residue fluctuations of RyR1 during MD simulation, we calculated the root mean square fluctuation (RMSF) as follows: first, we combined snapshots of a chosen ensemble; second, we superimposed the Cα coordinates onto the initial structure to attain a minimal root mean square deviation (RMSD); finally, we calculated the following RMSF at residue position *n*: 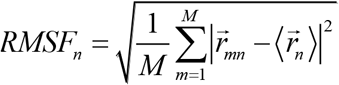, where 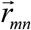 is the Cα position of residue *n* in snapshot *m*, 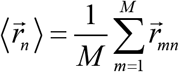 is the average Cα position of residue *n*, and *M* =1350 is the total number of snapshots (150 per trajectory from 9 trajectories).

### Analysis of inter-subunit contact surface area

We used the measure command of the VMD program (50) to calculate the solvent accessible surface area of each subunit in isolation (denoted ai for subunit i) and in a pairwise complex (denoted Aĳ between two adjacent subunits i and j), and then calculated the contact surface area between subunits i and j as (a_i_ + a_j_ - A_ij_) / 2.

### Hydrogen bond analysis

We used the following geometric criteria to identify a hydrogen bond (HB) between two polar non-hydrogen atoms (i.e., acceptor and donor): the donor-acceptor distance is < 3.5 Å, and the deviation of the donor-hydrogen-acceptor angle from 180° is < 60°. We used the VMD program (50) to identify and calculate the occupancy of each HB within a structural ensemble. For each HB-forming residue pair, we calculated its occupancy by adding the occupancies of HBs between them for all four subunits and then dividing it by 4. We only considered those HB-forming residue pairs with occupancy ≥ 0.3.

### Rg analysis

To probe inward/outward motions in RyR1 at the residue level of detail, we used the measure command of the VMD program (50) to calculate the radius of gyration Rg (n) from the Cα coordinates of four equivalent residue positions n of the RyR1 tetramer. Then we averaged Rg (n) over each MD-generated ensemble, and computed the fractional change of average Rg (n) from the Ca^2+^-unbound ensemble to the Ca^2+^-bound ensemble.

### PCA analysis

To identify dominant modes of motions/fluctuations sampled by MD simulation, we performed the principal component analysis (PCA) as follows: first, we combined snapshots of two ensembles (-Ca/Ca and +Ca/Ca); second, we superimposed the Cα coordinates of each snapshot onto the initial structure with a minimal RMSD; third, we calculated a co-variance matrix comprised of the following 3×3 block matrices 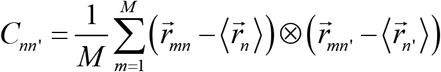 (see above for definitions of symbols); fourth, we diagonalized the co-variance matrix and kept the first and second dominant PCA modes which have the highest and second highest eigenvalues.

## Results and Discussion

### MD simulation of core RyR1 in the presence or absence of bound and solvent Ca^2+^

To understand how Ca^2+^ modulates the RyR1 structure and dynamics, we performed extensive MD simulation of the core RyR1 (see Fig 1) with or without Ca^2+^ bound, and in the presence of Ca^2+^ or K^+^ in solvent (see Methods). The reason for excluding the peripheral domains is because they were not resolved with sufficient resolution for all-atom MD simulation. Additionally, the peripheral domains are expected to play regulatory roles while the essential function of Ca^2+^ activation resides within the core RyR1 (22).

The initial RyR1 core structure was truncated from a 3.6-Å cryo-EM structure of rabbit RyR1 bound with Ca^2+^ only (PDB id: 5T15), which was embedded in a lipid bilayer and surrounded by a box of water molecules, Ca^2+^ or K^+^, and Cl^−^ ions (see Methods). We constructed the following three structural ensembles based on nine 200-ns MD trajectories per system: 1. Ca^2+^-bound RyR1 in the presence of Ca^2+^ and Cl^−^ in solvent (denoted +Ca/Ca); 2. Ca^2+^-unbound RyR1 in the presence of Ca^2+^ and Cl^−^ in solvent (denoted -Ca/Ca); 3. Ca^2+^-unbound RyR1 in the presence of K^+^ and Cl^−^ in solvent (denoted -Ca/K). A comparison between system 1 and 2 will reveal the effects of Ca^2+^ binding (to the activating site). A comparison between system 2 and 3 will reveal the effects of solvent Ca^2+^.

To assess the stability of MD simulation, we calculated the root mean square deviation (RMSD) of Cα atoms relative to the initial structure for each trajectory (see Fig S1). While most trajectories exhibited saturation of RMSD after < 50 ns, we observed relatively large RMSD values (4-8 Å, see Fig S1) which may be attributed to large intrinsic inter-subunit motions/fluctuations required for channel opening. Indeed, we observed similarly large RMSD in our previous MD simulation of the tetramer of RyR1 N-terminal domains (25). Alternatively, the large RMSD may be attributed to missing peripheral domains that destabilize the core structure. This possibility will be addressed by future MD simulation when new RyR1 structures with better-resolved peripheral domains become available.

Among the three ensembles, we observed least variations in RMSD in -Ca/Ca (see Fig S1), hinting for higher stability and lower flexibility characteristic of the closed state (18). We attributed this to strong binding of solvent Ca^2+^ to the RyR1 core that stabilizes a more compact RyR1 in the closed state (e.g. via neutralization of the negative charges). Indeed, the RyR1 core is rich in negative charges (with 182 acidic residues and 101 basic residues per subunit). In contrast, RyR2 has fewer negative charges (with 166 acidic residues and 111 basic residues per subunit), resulting in weaker binding to solvent Ca^2+^. Indeed, RyR2 is activated by Ca^2+^ to a greater extent and requires higher Ca^2+^ concentrations for inhibition than RyR1 (51). Additionally, RyR1/RyR2 chimeras containing an RyR2 C-terminal core (residues 3726–5038) showed reduced channel inhibition at elevated Ca^2+^ levels, suggesting that the RyR1/RyR2 core contains inhibitory Ca^2+^ binding sites with different affinities (52). Further supporting the importance of electrostatic interaction to inhibition, other di-valent cations like Mg^2+^ (53) and Zn^2+^ (54) also inhibit RyRs at millimolar concentrations (55).

To locate possible Ca^2+^ binding sites, we counted the number of solvent Ca^2+^ adjacent to each residue position (within a 5-Å cutoff distance), and found that Ca^2+^ is densely populated near clusters of acidic residues in the following regions: EFH12 (residues 4116 - 4119), luminal loop (residues 4867 - 4870), and other surface loops (residues 3684 - 3691, 3740 - 3750, and 4584 - 4625) (see Fig S2). We hypothesize that Ca^2+^ binding to some of these sites may inhibit RyR1 activity. Consistent with our hypothesis, previous studies have located possible Ca^2+^ binding sites overlapping with the above regions [residues 3657 - 3776 (56), 4381 - 4626 (8), and two EF-hand motifs (6, 57)]. Two regions of RyR2 (residues 4020 - 4250 and 4560 - 4618) were implicated in Ca^2+^-dependent inactivation (58). Another study identified a Ca^2+^ inhibition site between residues 4063 and 4209 in RyR1 (59). Our finding will guide future mutational studies to precisely pinpoint the binding sites for Ca^2+^ inhibition.

### RMSF analysis found dual Ca^2+^-dependent modulation of RyR1 flexibility

To analyze the flexibility of core RyR1 at the residue level of resolution, we calculated the RMSF profile for each of the three ensembles (sees Methods), which measures the average atomic fluctuations of each residue during MD simulation. While the RMSF profiles of +Ca/Ca and −Ca/Ca are similar (see Fig 2a), Ca^2+^ binding slightly increases RMSF in the TMD (see the higher local minima of RMSF in +Ca/Ca in Fig 2a). In contrast, the RMSF profiles of −Ca/Ca and −Ca/K are notably distinct, with the solvent Ca^2+^ reducing RMSF in the cytosolic domains and the pore domain in −Ca/Ca (see Fig 2b). Therefore, while the solvent Ca^2+^ reduced RyR1 flexibility, Ca^2+^ binding (at the activating site) increased RyR1 flexibility when Ca^2+^ is present in solvent. Such dual modulation effects of Ca^2+^ on RyR1 flexibility support the possibility for Ca^2+^ to both activate and inhibit RyR1 activity.

**Figure 2.**
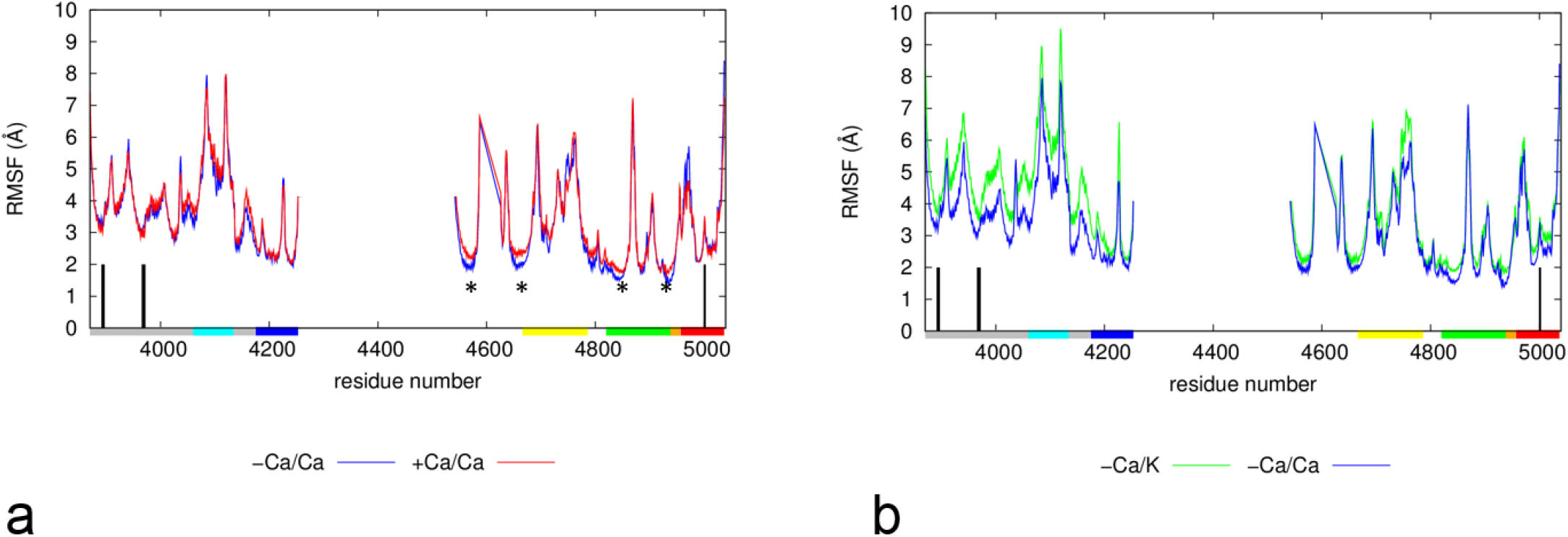
The Ca^2+^-dependent RMSF profiles of RyR1 core as compared (a) between +Ca/Ca and -Ca/Ca, (b) between -Ca/K and -Ca/Ca. The key domains are marked by bars on the horizontal axis colored as follows: CTD (red), TAF (blue), S6C (orange), pore domain (green), S2S3 (yellow), and EFH12 (cyan). The positions of Ca^2+^-binding residues (E3893 E3967 Q3970, H3895, and T5001) are marked by black vertical lines. In panel a, the locations of RMSF differences are marked by *.

Among various RyR1 domains, EFH12 has the highest RMSF followed by S2S3 and CTD, while TAF and the pore domain have low RMSF except in their solvent-exposed regions (see Fig 2). At the activating Ca^2+^-binding site between CSOL and CTD, residues E3893, E3967, Q3970, and H3895 of CSOL have low RMSF, while T5001 of CTD exhibits a peak in RMSF (see Fig 2), which indicates a dynamic CTD moving relative to a static CSOL. The high flexibility of EFH12, S2S3, and CTD is consistent with the hypothesis that these domains may undergo allosteric motions to couple Ca^2+^ binding to the pore opening.

### The inter-subunit interfaces of RyR1 were oppositely modulated by bound and solvent Ca^2+^

In our previous MD simulation of the tetramer of RyR1 N-terminal domains (25), we found that many disease mutations perturb the inter-subunit interactions, thus destabilizing RyR1 so it opens more readily. To test if Ca^2+^ effects the inter-subunit interfaces in the core RyR1, we analyzed the flexibility of inter-subunit interfaces by calculating the following three quantities: 1. inter-subunit distances (i.e. the distance between the centers of mass of two adjacent RyR1 subunits); 2. inter-subunit contact surface area (see Methods); 3. number of inter-subunit hydrogen bonds (see Methods). The findings are summarized as follows (see Fig 3):

In comparison with −Ca/K, the presence of solvent Ca^2+^ in +Ca/Ca and −Ca/Ca shifts the distributions of inter-subunit distance to the left (see Fig 3a), resulting in a more compact tetramer consistent with a closed/inhibited RyR1 channel.

**Figure 3.**
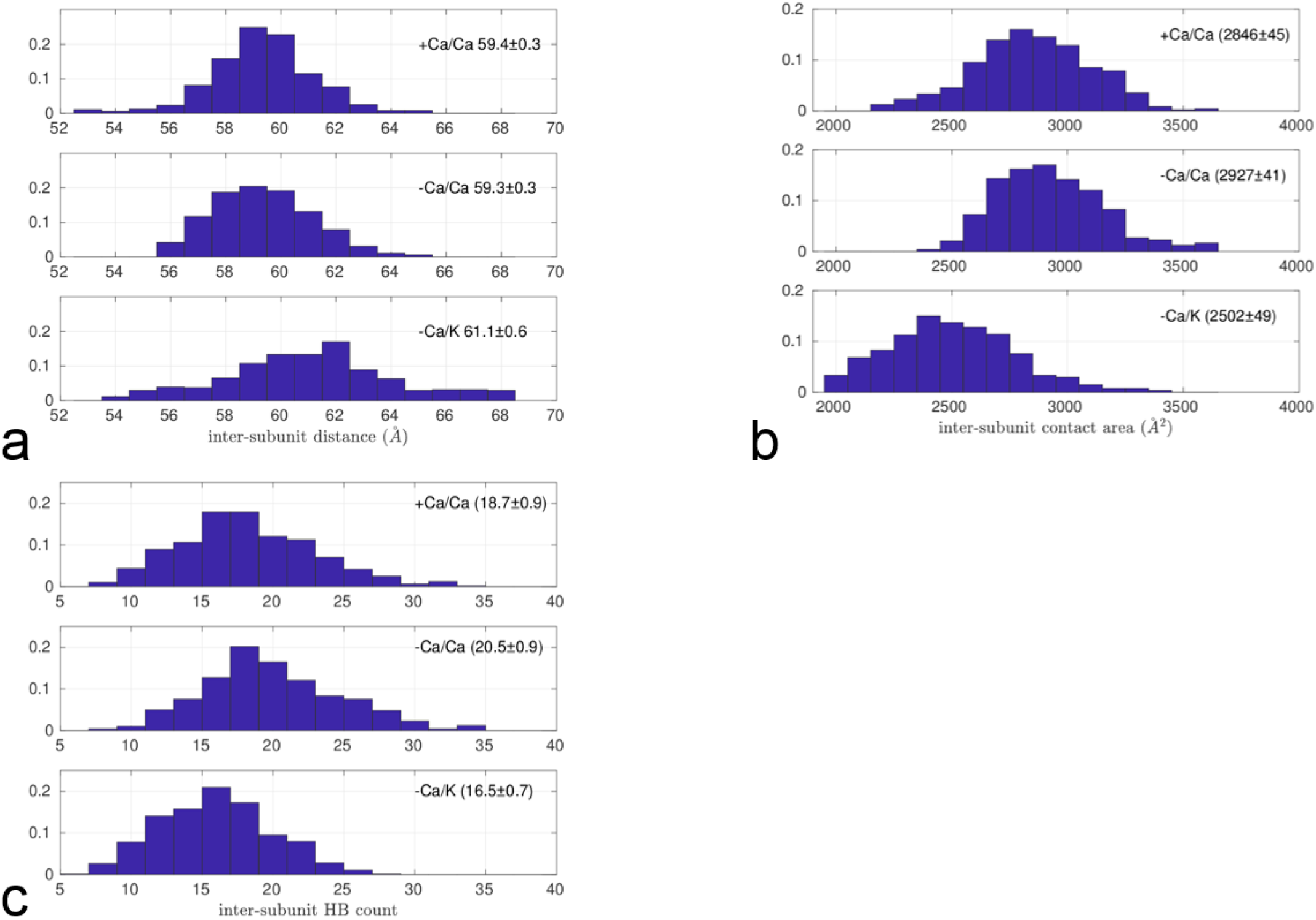
Histograms of (a) inter-subunit distance, (b) inter-subunit contact surface area, (c) number of inter-subunit HBs for three structural ensembles of RyR1 core (+Ca/Ca, -Ca/Ca, and -Ca/K). Mean ± standard error are shown in each histogram.

In comparison with −Ca/K, the presence of solvent Ca^2+^ in +Ca/Ca and −Ca/Ca shifts the distributions of inter-subunit surface area and hydrogen bonds to the right, resulting in a more stable and inhibited RyR1 tetramer. In comparison with −Ca/Ca, Ca^2+^ binding in +Ca/Ca significantly decreases the inter-subunit contact surface area and the number of inter-subunit hydrogen bonds (see Fig 3b, 3c), leading to a less stable and more active RyR1.

Therefore, solvent Ca^2+^ and Ca^2+^ binding (at the activating site) have opposite effects on the inter-subunit interfaces, with the latter weakening while the former strengthening inter-subunit interactions, resulting in greater and lower flexibility, respectively. This is consistent with the above finding of dual effects of Ca^2+^ on RMSF.

### Rg analysis revealed Ca^2+^-dependent expansion/contraction of RyR1

Although cryo-EM has resolved the open structures of RyR1 in complex with Ca^2+^/ATP/Caffeine (22), it is unknown how RyR1 is activated via conformational changes triggered by Ca^2+^ binding alone. Toward answering this question, we have analyzed the conformational shifts of the RyR1 core upon Ca^2+^ binding (at the activating site) or in the presence of solvent Ca^2+^. For simplicity, we focused on those symmetric inward/outward domain motions relevant to gating (e.g., channel pore closing/opening). To this end, we calculated ensemble averages of the radius of gyration (Rg) at each residue position of the core RyR1 (see Methods). An increase/decrease in Rg would indicate an outward/inward domain motion that could be coupled to activation/inhibition of RyR1. This analysis was previously applied to the heat-activated gating of the TRPV1 channel (60).

To show the effectiveness of the Rg analysis, we calculated the changes in Rg from the Ca^2+^-bound structure (PDB id: 5T15) to the apo closed structure (PDB id: 5TB0), and to the Ca^2+^/ATP/Caffeine-bound open structure (PDB id: 5T9V). As expected, the Rg changes to the open structure are more pronounced than those to the closed structure, indicating large expansion in S6C and CTD, and less expansion in S4S5L, TAF, and S2S3 (see Fig 4a). These observed domain motions are consistent with several possible allosteric pathways connecting the Ca^2+^-binding site (at the CTD/CSOL interface) to the channel pore (e.g., CTD ➔ S6C, CSOL ➔ EFH12 ➔ S2S3 ➔ S4S5L ➔ S6C, see Fig 1c).

Next, to probe the activating effect of Ca^2+^ binding, we calculated the average Rg changes from the -Ca/Ca ensemble to the +Ca/Ca ensemble in the presence of solvent Ca^2+^. We found small but significant Rg increases in EFH12, TAF, S2S3, and CTD (but not in S4S5L, see Fig 4b), which is qualitatively similar to the cryo-EM-observed changes (see Fig 4a). In the pore domain, we observed Rg increases in S6C and the selectivity filter (residues 4894 - 4900) in agreement with a cryo-EM study (23) (see Fig 4b). To study the inhibitory effect of solvent Ca^2+^, we analyzed average Rg changes from the -Ca/K ensemble to the -Ca/Ca ensemble (see Fig 4c), and found Rg decreasing extensively, which is consistent with channel contraction indicative of inhibition.

**Figure 4.**
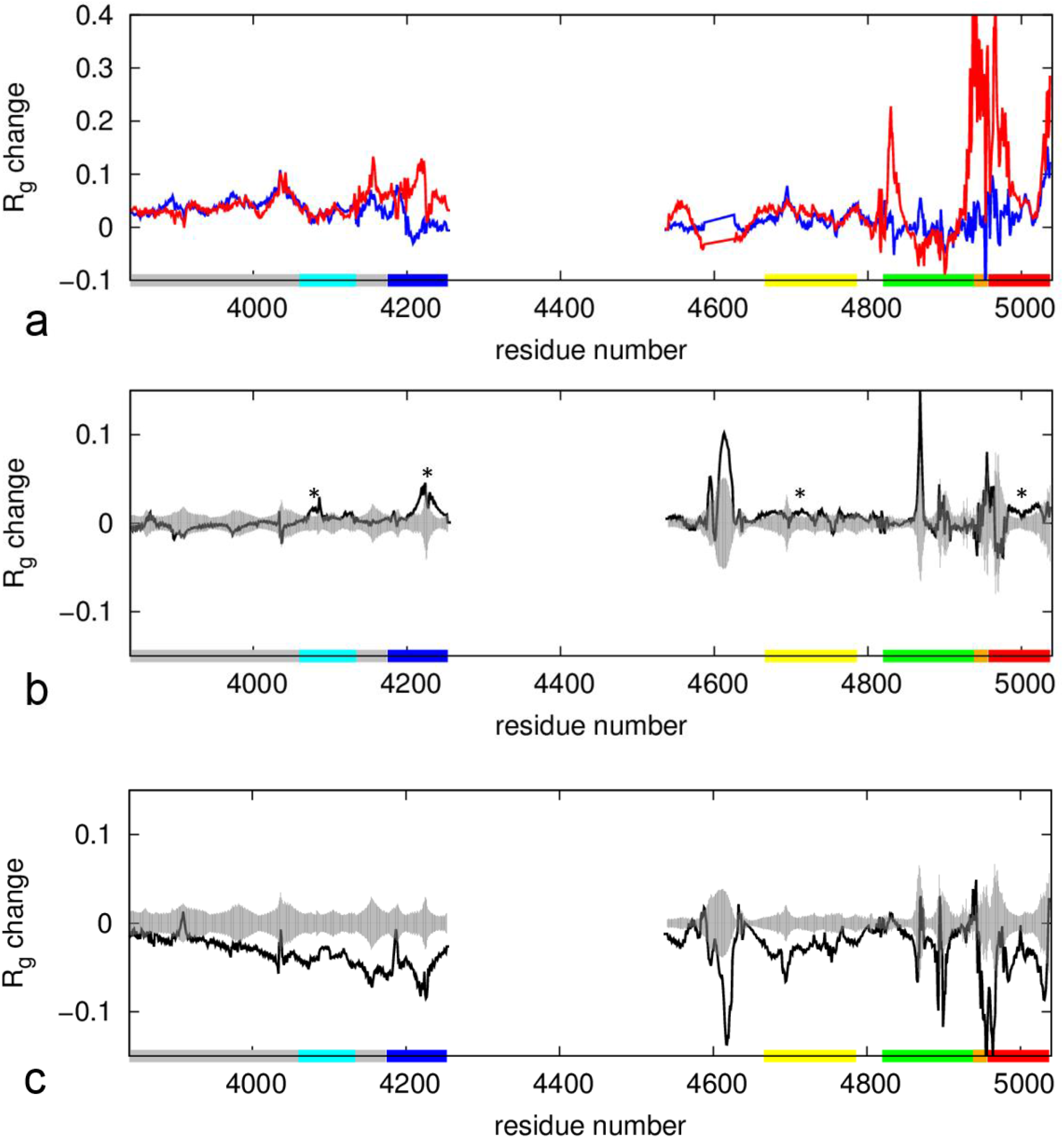
Fractional changes of Rg as a function of residue positions: (a) from the Ca^2+^-bound cryo-EM structure to the open structure (red) and to the closed structure (blue); (b) from the −Ca/Ca ensemble to the +Ca/Ca ensemble; (c) from the −Ca/K ensemble to the −Ca/Ca ensemble. Panel b and c show the ensemble-averaged Rg changes (black curves) and corresponding standard errors (gray shades). The key domains are marked by bars on the horizontal axis colored as follows: CTD (red), TAF (blue), S6C (orange), pore domain (green), S2S3 (yellow), and EFH12 (cyan). In panel b, the locations of signficant Rg differences are marked by *.

In sum, we observed small but significant expansion in key domains, including EFH12, TAF, S2S3, and CTD, after Ca^2+^ binding (to the activating site), which may be coupled to downstream motions in the pore domain for activation. In contrast, the presence of solvent Ca^2+^ resulted in extensive contraction of the RyR1 core characteristic of inhibition. This is consistent with the above findings of dual effects of Ca^2+^ on flexibility and intersubunit interactions.

### Channel pore analysis reveals Ca^2+^-dependent partial pore expansion

To assess the open/closed state of the RyR1 channel pore and its dynamics during MD simulation, we used the HOLE program (61) to calculate the pore radius along the pore axis for each of the three ensembles. Similar to the Ca^2+^-bound cryo-EM structure (22), the pore remains narrowly constricted near I4937 (with pore radius ~ 1 Å) consistent with a closed channel. However, upon Ca^2+^ binding, the pore shows modest expansion in the selectivity filter (near G4894) and S6C (near T4956) (see Fig S4a), suggesting that RyR1 was primed for activation. In contrast, solvent Ca^2+^ caused the upper and the lower pore to contract (see Fig S3b), suggesting inhibition. Additionally, the upper pore (in S6C and CTD) is more dynamic than the lower pore as indicated by the larger variations in pore radius (see Fig S4). This is consistent with the above findings of opposite effects of solvent and bound Ca^2+^, which are inhibitory and activating, respectively.

### PCA analysis revealed Ca^2+^-activated asymmetric inter-subunit opening/closing motions

To further analyze how the structural fluctuations of core RyR1 are modulated by Ca^2+^, we applied the principal component analysis (PCA) to the MD-generated ensembles (see Methods). The PCA aims to identify the dominant modes of structural fluctuations/motions as sampled by MD simulation (for the +Ca/Ca and −Ca/Ca systems). Given the four-fold symmetry of RyR1 homo-tetramer, we divided all PCA modes into two groups (symmetric modes that satisfy the four-fold symmetry, and asymmetric modes that violate this symmetry), and then selected the dominant mode from each group for further analysis.

The first mode (PC1) accounts for 11% of the total conformational variations, and it depicts asymmetric outward/inward motions of two diagonally opposite subunits (Fig 5b), which are apparently coupled to asymmetric opening of the channel (i.e. via pulling of the two outward-moving subunits). The second mode (PC2) accounts for 7% of all variations, and it captures distinct swiveling/twisting motions of individual subunits (Fig 5c), which are not apparently coupled to channel opening/closing. Both modes exhibit high asymmetry, in contrast to the cryo-EM RyR1 structures that obey four-fold symmetry. Interestingly, we observed asymmetric channel opening in TRPV1 (a tetrameric channel with a similar 6TM-fold of TMD) (60).

**Figure 5.**
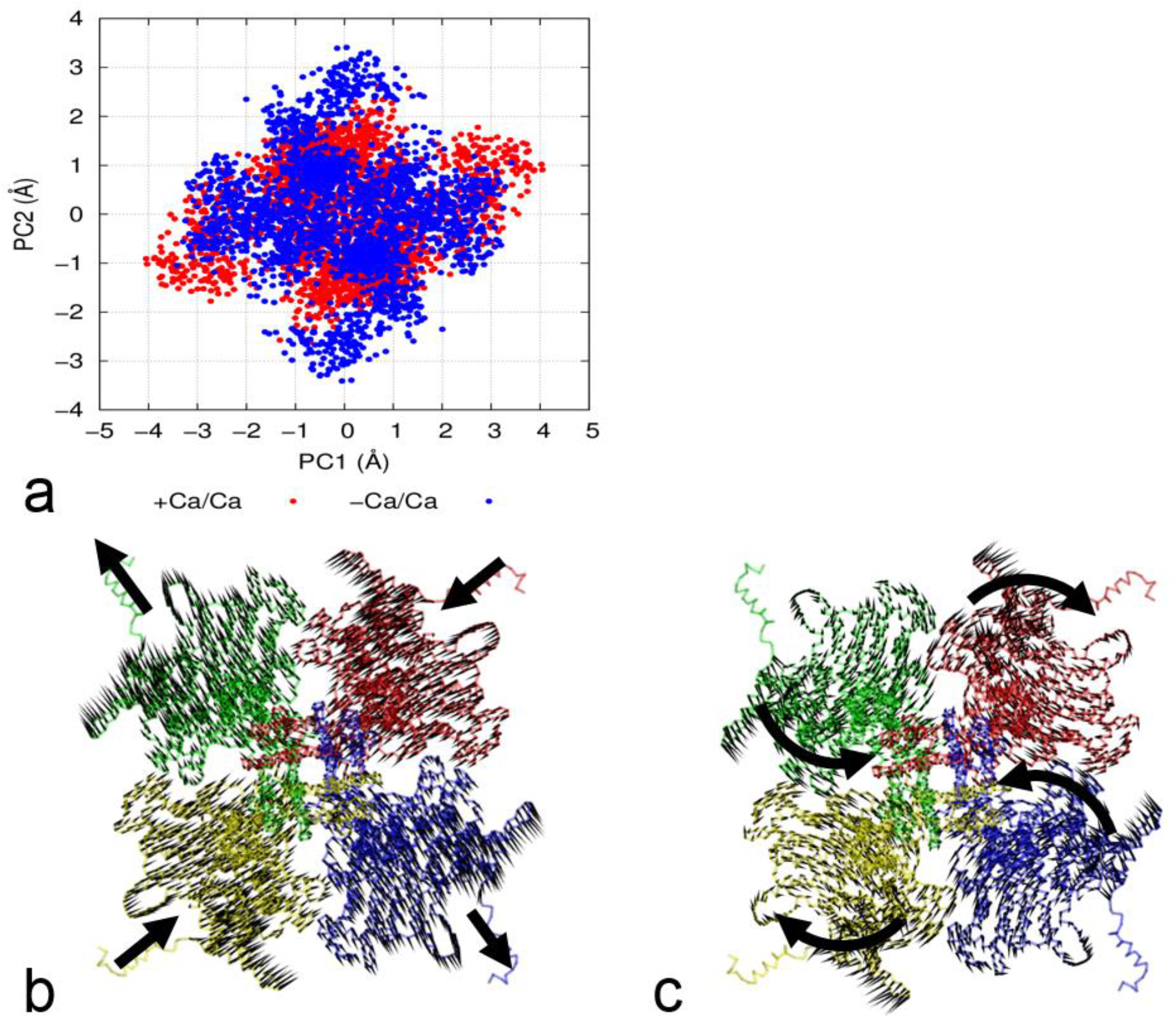
Results of principal component analysis: (a) projections of the −Ca/Ca ensemble (blue) and the +Ca/Ca ensemble (red) onto a plane spanned by the first and second dominant PCA modes (PC1 and PC2). (b) the vector field plot for the motions of individual residues as depicted by the first dominant mode (PC1). (c) the vector field plot for the second dominant mode (PC2). Panels (b) and (c) use the same bottom view as in Fig 1b with four subunts colored distinctly and their motions indicated by bold arrows.

To probe how Ca^2+^ binding modulates the structural fluctuations of the RyR1 core along the above two dominant modes, we projected snapshots of the two ensembles (+Ca/Ca and −Ca/Ca) onto a plane spanned by the eigenvectors of these two modes (see Fig 5a). While the two ensembles largely overlap, +Ca/Ca is more spread out along the first mode while -Ca/Ca is more extended along the second mode (see Fig 5a), suggesting that Ca^2+^ binding selectively enhances the first mode while suppressing the second mode. Therefore, we propose that Ca^2+^ binding favors RyR1 activation by amplifying the gating-relevant opening/closing mode (PC1) while suppressing the other modes which are less relevant to gating (e.g. PC2).

### Disease mutation sites mediate allosteric couplings via hydrogen bonds formed at inter-domain interfaces

Many disease mutations have been identified in RyR1, most of which cause gain of function that results in a leaky channel. However, it remains unknown how they alter the RyR1 activity at the molecular level. Based on our previous and present simulations, we hypothesize that these mutations perturb functionally important interactions that stabilize or couple various RyR1 domains to maintain a balanced equilibrium between the closed state and the open state. To substantiate this hypothesis, we examined hydrogen bonds (HBs) dynamically formed during MD simulation that involve known disease mutations of human RyR1 obtained from the HGMD database (http://www.hgmd.cf.ac.uk). Total 70 mutations were matched with high-occupancy HBs formed in either the −Ca/Ca or the +Ca/Ca ensemble (see Table S1). Many of them were formed within each domain in a Ca^2+^-independent manner, so they likely stabilize the domain structure. Other mutation sites were located at the interfaces between key domains of the RyR1 core structure (including EFH12, TAF, S2S3, S4S5L, S6C, and CTD, see Fig 1), which will be the focus of our following analysis. Here each HB is denoted by a pair of donor/acceptor residues (using rat RyR1 residue numbers), while the position of each disease mutation is represented using human RyR1 residue numbers (see Table S1). Our discussion is organized by individual domains where the mutations are located.

#### ♦ TAF

Three mutations in TAF (R4179H, E4181K, and A4185T) are involved in multiple HBs (R4180 - E4981, R4180 - E4982, R4180 - N4987, E4182 - Y4988, and A4186 - Q5006) that couple TAF to CTD. Another mutation (E4243K) perturbs an HB (E4244 - K4665) that couples TAF to S2S3. Although the total number of HB changes little between -Ca/Ca (53.0±2.3) and +Ca/Ca (54.8±2.5), the above HBs between TAF and CTD have higher occupancy in −Ca/Ca (see Table S1), and their vdW interaction is also stronger in −Ca/Ca (−653.9±8.1 kcal/mol) than in +Ca/Ca (−643.0±6.4 kcal/mol), which are consistent with a more dynamic CTD (relative to TAF) in the +Ca/Ca ensemble. In contrast, the TAF-S2S3 interactions change little between −Ca/Ca and +Ca/Ca, with a vdW energy of −81.9±4.3 kcal/mol in −Ca/Ca and −85.2±3.3 kcal/mol in +Ca/Ca.

#### ♦ S2S3

Two mutations in S2S3 (R4737Q and R4737W) involve two inter-subunit HBs (D4079 - R4736 and E4075 - R4736) that couple S2S3 to EFH12, and one of them (E4075 - R4736) only forms in −Ca/Ca. Consistent with a stronger S2S3-EFH12 coupling in −Ca/Ca, we observed a stronger S2S3-EFH12 vdW interaction in −Ca/Ca (−64.0±4.4 kcal/mol) than in +Ca/Ca (−54.8±6.3 kcal/mol), although the total numbers of HBs are only slightly different (13.1±1.1 in −Ca/Ca and 12.2± 1.3 in +Ca/Ca). Interestingly, the EFH12 – S2S3 contacts were formed in the Ca^2+^-only structure, but were broken in the open structures (see ref (22)).

#### ♦ S4S5L

A mutation in S4S5L (T4823M) perturbs an HB (E4212 - R4824) that couples S4S5L to TAF, with lower occupancy in +Ca/Ca than in −Ca/Ca, suggesting a more flexible S4S5L-TAF interface in +Ca/Ca. Consistent with this, we observed a stronger TAF - pore domain vdW interaction in −Ca/Ca (−80.6±2.7 kcal/mol) than in +Ca/Ca (−74.0±3.9 kcal/mol), although the total numbers of HBs are only slightly different (12.4±1.4 in +Ca/Ca and 11.0± 1.5 in −Ca/Ca).

#### ♦ S6C

One mutation in S6C (L4936R) involves an HB (N4833 - L4935) that couples S6C to S4S5L. Another mutation (D4939E) involves an inter-subunit HB (D4938 - R4944) that couples S6C to S6C’ of an adjacent subunit. While the first HB changes little between −Ca/Ca and +Ca/Ca, the second has a higher occupancy in +Ca/Ca. This finding agrees with the cryo-EM observation that R4944 of S6C shifts from interacting with E4942 on S6C’ in the closed structure to interacting with D4938 on S6C’ in the open structure (see ref (22)).

#### ♦ CTD

One mutation in CTD (G5006S) perturbs an HB (Q3970 - G5005) coupling CTD to the Ca^2+^-binding site in CSOL, which is only present with high occupancy in +Ca/Ca, suggesting an enhanced CSOL-CTD coupling upon Ca^2+^ binding. Consistent with this, we observed a stronger CSOL-CTD vdW interaction in +Ca/Ca (−122.9±4.6 kcal/mol) than in -Ca/Ca (−113.6±8.2 kcal/mol), although the total numbers of HBs are unchanged (11.2±1.5 in +Ca/Ca and 11.1±1.4 in −Ca/Ca).

In summary, by correlating known disease mutation sites with MD-observed inter-domain/subunit interactions, we have identified Ca^2+^-dependent couplings (via vdW interactions and specific HBs) for the following domain pairs: CSOL-CTD, CTD-TAF, EFH12-S2S3, and TAF-S4S5L. Along with the covalent coupling of CTD to S6C, the above findings are most consistent with the CSOL ➔ CTD ➔ S6C pathway for Ca^2+^-activation. An alternative pathway of CSOL ➔ CTD ➔ TAF ➔ S4S5L ➔ S6C is also consistent except for the S4S5L-S6C coupling not showing Ca^2+^-dependent changes, possibly owning to short MD simulation. Another possible pathway is via EFH12 ➔ S2S3 ➔ S4S5L, although our simulation did not find direct coupling from S2S3 to S4S5L.

## Concluding remarks

To elucidate how Ca^2+^ modulates RyR1 structurally and dynamically, we performed extensive MD simulation of the core RyR1 structure in the presence or absence of bound and solvent Ca^2+^. In the presence of solvent Ca^2+^, Ca^2+^ binding to the activating site resulted in elevated dynamics of RyR1 embodied by higher inter-subunit flexibility, asymmetric inter-subunit motions, outward domain motions and partial pore dilation, which may initiate early events that lead to subsequent channel opening. In contrast, the presence of solvent Ca^2+^alone caused RyR1 contraction with reduced dynamics. Therefore, Ca^2+^ can modulate RyR1 dynamics both positively and negatively, presumably by binding to different sites. Based on the simulation, we propose that Ca^2+^ binding (at the activating site) activates RyR1 by increasing its flexibility and destabilizing its inter-subunit interfaces. However, solvent Ca^2+^ inhibits RyR1 by strongly interacting with the negatively charged RyR1. Our simulation has revealed solvent Ca^2+^ densely distributed near clusters of acidic residues (see Fig S2), which will guide future site-directed mutagenesis studies to precisely pinpoint the binding sites for Ca^2+^ inhibition. Alternatively, the inhibitory effect of solvent Ca^2+^ is via nonspecific electrostatic interactions with a large number of acidic residues in RyR1, which will need to be probed by chimera-based studies (52)..

It is well-known that the open probability (Po) of RyR1 has a biphasic dependence on cytosolic Ca^2+^ concentration, with a maximal Po in the 10-to 100-mM range and lower activity at both higher and lower levels of Ca^2+^ (62). Our finding of opposite effects of bound Ca^2+^ and solvent Ca^2+^ on RyR1 structure and dynamics provided some mechanistic insights to this: if the Ca^2+^ concentration is too low, Ca^2+^ cannot bind to the activating site to enhance the RyR1 dynamics, resulting in low activity; if the Ca^2+^ concentration is too high, the inhibitory effect of solvent Ca^2+^ would dominate the activating effect of Ca^2+^ binding (at the activating site), resulting in low activity; only at intermediate Ca^2+^ concentration, Ca^2+^ binding can effectively activate RyR1 despite the inhibitory effect of solvent Ca^2+^.

Combining our simulations with the map of disease mutation sites in RyR1, we constructed a wiring diagram of key functional domains linked by specific hydrogen bonds involving the disease mutation sites, which hint for possible allosteric coupling pathways for Ca^2+^ activation. Two domains (CTD and TAF) stand out to be critically involved in Ca^2+^-dependent inter-domain interactions (see Fig 1c), supporting their roles in transmitting the signal of Ca^2+^ binding to the pore domain (via S6C and S4S5L, see Fig 1c). In addition to allosteric couplings, many mutation sites involve HBs that stabilize individual domain structures and their interfaces (see Table S1). The mutational perturbation to these stabilizing interactions can shift the equilibrium from the more stable closed state toward the less stable open state, resulting in an overactive and leaky RyR1 channel (63).

Owning to the short simulation time, the MD-observed Ca^2+^-dependent conformational changes are smaller and less extensive than the full-blown changes to the open structure as observed by cryo-EM (22). Some key structural changes in the pore domain were not observed, such as S4S5L straightening and S6C opening (22). For future work, we will extend our MD simulation to multi-microseconds to better explore the activation transition triggered by Ca^2+^ binding.

## Author Contributions

WZ designed research; WZ and HW performed research; WZ and HW analyzed data; WZ wrote the paper.

## Acknowledgement

This study was supported by a grant from American Heart Association (#17GRNT33690009). The simulations were conducted using the supercomputing cluster of the Center for Computational Research at the University at Buffalo.

